# How drought affects range and variability of environmental conditions in subtropical South African estuaries

**DOI:** 10.1101/2022.02.17.480947

**Authors:** UM Scharler, SJ Bownes, H Jerling

**Author notes:** ^#^Corresponding author (US). Current address: Department of Biology and Environmental Sciences, University of Mpumalanga, South Africa (SJ Bownes).

## Abstract

Globally estuaries are under increasing pressure from human development impacts as well as the growing effects of climate change that is already, and increasingly so, causing major changes in events such as droughts. Although droughts are regular occurrences and are projected to become progressively more significant in many parts of the world, their potential impacts on estuaries requires better understanding and quantification. This study aims to quantify environmental and phytoplankton parameter changes in two contrasting subtropical estuary types in South Africa; the temporarily closed uMdloti Estuary and the predominantly open to the sea uMlalazi Estuary during a hydrological drought event and subsequent non-drought period. Drought caused lower salinities, almost exclusively freshwater, throughout the temporarily closed estuarine type uMdloti. The marine-estuarine salinity environment of the predominantly open uMlalazi during drought expanded to include lower salinities in the non-drought period. Dissolved oxygen levels were more variable during drought conditions including periods of hypoxia, but more stable at elevated levels in both estuaries during the non-drought period. Temperature measurements included higher values during drought, especially in the dry season. Chl-a concentrations were overall higher and more variable during drought in uMdloti, with periods of bloom formation as well as decay. Drought-induced conditions can span several generations for estuarine and estuarine associated organisms. The results from the study provide quantitative support for the hypothesis that extended periods of drought conditions lead to a decrease in niche availability to estuarine organisms in a range of estuary types.

## Introduction

The importance of estuaries for biodiversity, productivity and ecosystem services to humans is undisputed. Evaluations of drought impacts on estuarine environments are therefore important for several reasons. Estuaries exhibit a high variability in environmental conditions arising from their connection to both the sea and rivers, and thus they harbour distinct biodiversities and communities that are adapted to such conditions (Elliott and Quintino, 2007). They furthermore provide habitat to marine organisms that use estuaries as nurseries including commercially important fish and invertebrate species (de Villiers and Hodgson, 1999; Lamberth et al. 2009; Harrison and Whitfield, 2021). They improve the water quality of riverine inflow, and feature high productivity. All aspects together make them valuable to human society as they support food production, improve pollution levels and provide places of recreation (Barbier et al. 2011). Changes to the estuarine environment can disrupt such services, and we aimed to document how drought affects specifically which aspects of the estuarine environment. IPCC (2021) projections for the South African region are towards more intense ecological and agricultural droughts, which have an effect on freshwater abstractions from the catchment, and thus riverflow and freshwater input into estuaries. Even though South African legislation requires environmental flows to be set to achieve desired ecological states (National Water Act (Act No. 36 of 1998)), this may not be feasible, and is often not adhered to, under frequent or prolonged drought conditions and therefore it is important to know what type of environment future estuaries may offer to its biota. As climate and weather change projections increase their global accuracy and mitigation efforts are gearing up, human society needs more information on possible consequences of climate change. This is important not only for land-based food production (Conway et al., 2015), but also for the coastal and marine food supply and the biodiversity that underpins it in particular geographic and climatic regions.

Estuarine biota are able to cope with its intrinsically variable environment either by tolerating wide ranges of physico-chemical parameters, or by exploiting confined physico-chemical niches within the estuary. For instance, the degree of tolerance to salinity ranges prompted a categorisation into eury- or stenohaline estuarine taxa (Smyth and Elliott, 2016). Temperature influences the metabolic activity and reproductive cycles of estuarine organisms (Moens and Vincx, 2000; Yvon-Durocher et al., 2012; Tagliarolo et al., 2018) and thereby it is an important aspect of estuarine ecology. While many estuarine taxa have wide temperature tolerances, the optimal and critical thresholds for performance can vary between species and life stages (Jeffries et al., 2016; Payne et al., 2016). Temperature in estuaries varies as a combination of riverine, oceanic and atmospheric temperature fluctuations and changing temperatures have received increased attention within the context of climate change and biogeographic range shifts (Jeffries et al., 2016; Scanes et al., 2020; Whitfield et al., 2016).

Vertical and longitudinal gradients of dissolved oxygen (DO) concentrations provide niches that organisms with different preferences can exploit. Hypoxia and anoxia is a known agent of fish kills (Wong et al. 2008), and hypoxic stress for benthic organisms increases both the mortality rate and the threshold DO concentrations at which mortality increases (Vaquer-Sunyer and Duarte, 2011) even that some estuarine benthic animals are actually adapted to short periods of hypoxia (Sagasti et al., 2000). Tightly coupled with changes in DO concentrations are those of pH. During autotroph bloom conditions, the use of CO_2_ and increase of oxygen production by primary producers such as phytoplankton and macroalgae increases pH. This bloom condition is followed by decay and subsequent lowering of DO levels and pH (Carstensen and Duarte, 2019). Not as prominent a topic in estuaries than in the ocean until about a decade ago, changes in pH as a result of rising pCO_2_ is studied increasingly in estuaries (Feely et al., 2010; Scanes et al., 2020). Wide ranging impacts of changing pH levels can have effects on population, community and ecosystem levels, and may act in conjunction with temperature to influence metabolism, or predator-prey relationships through changes in prey vulnerability (e.g. calcifiers) or species performance (Gaylord et al., 2015).

In short, estuarine biotic communities reflect their environmental conditions, and undergo biodiversity and community changes as a result of it. The natural spatial and temporal variations in estuarine environments therefore support the biodiversity, productivity and functioning of estuaries (Elliott and Whitfield 2011). Pertinent from the above examples is that climate change and extreme climatic events (e.g. droughts and floods) are likely to have dramatic effects on estuarine environments. Community changes during drought conditions as a result of changing environmental parameters have been documented worldwide, especially from estuaries in Australia, South Africa, Brazil, USA or Portugal (e.g. Hastie and Smith, 2006; Dolbeth et al., 2010; Wetz et al., 2011; Carrasco and Perissinotto, 2012; Barroso et al., 2018). However, there are still fundamental gaps in our understanding of the effects of extreme and prolonged events on the physical, chemical and biological characteristics of estuaries, and consequently how estuarine water quality, ecological dynamics and ecosystems respond (Wetz and Yoskowitz, 2013; van Niekerk et al., 2019).

Seasonal rainfall and natural meteorological dry/wet cycles have a profound effect on riverflow and consequently on estuarine environmental conditions (Wolanski and Elliott 2015). There is already a tendency for intense and extensive droughts in South Africa that occur during periods of lower rainfall that last several years (Malherbe et al., 2016). Rainfall in South Africa varies on interannual (2-8 years), quasi-decadal (8-13 years) and inter-decadal (15-28 years) timescales (Malherbe et al., 2016; Pohl et al., 2018; Dieppois et al., 2019). Pertinent for our study, we experienced a drought period with extremely dry years during 2014-2016, and a non-drought period of average rainfall during 2019-2020 (Ndlovu and Demlie, 2020; South African Weather Service, 2020).

South Africa features nearly 300 estuaries of nine different types, and its most prominent type (> 70% of estuaries), are temporarily closed off from the sea under natural conditions (van Niekerk et al. 2020). Of the world’s such small, microtidal estuaries prone to closure, 16 % are situated in South Africa (McSweeney et al., 2017). They are closed off from the sea via a sand bar in seasons of lower rainfall and open with different frequencies during periods of higher rainfall. Under closed mouth conditions, small estuaries are more prone to riverine influence with concomitant shifts in the physico-chemical environment, community assemblages and productivity (Froneman, 2004; Nozais et al., 2005; Scharler et al., 2020). Many of the worldwide studies during drought conditions, including those from South Africa’s cold and warm temperate biogeographic regions, report hypersalinities in open and temporarily closed estuaries (Perissinotto et al. 2012; Hallett et al. 2017; Warwick et al. 2018). To the contrary, in South Africa’s subtropical region most of the temporarily closed estuaries are perched, with a water level higher than mean sea level when closed, and owing to minimal but constant freshwater input such estuaries freshen during closure (Perissinotto et al. 2012; Scharler et al. 2020).

The second most common type in South Africa is that of predominantly open estuaries which feature an open connection to the sea throughout the year, although some may close during prolonged periods of drought (Van Niekerk et al., 2019). This type can be prone to increased marinisation when open, depending on the amount of riverine input (Scharler and Baird, 2003). Here we aim to quantify spatial and temporal changes in environmental conditions and in microalgae chlorophyll-a concentrations under drought and non-drought conditions, with a focus on ranges and variability during dry and wet seasons.

## Material and methods

The study was conducted over a total period of seven years (2014-2020) spanning drought and non-drought conditions along the subtropical east coast of South Africa, which is a summer rainfall zone and receives some of the highest rainfall in South Africa (Dieppois et al., 2016; Roffe et al., 2021). The study sites encompass a large temporarily closed estuary (LTCE) as well as an estuary that is predominantly open to the sea (PrOE) and only closes during major prolonged droughts (Figure 1; van Niekerk et al., 2019). The LTCE uMdloti (31°7.’44.9328”, 29°39’2.1348”) is situated within the eThekwini Municipality, north of Durban. Riverine flow reaching the estuary is modified in terms of quantity and quality. The catchment area of 486 km^2^ delivers a mean annual runoff of 71.9 Mm^3^ year^−1^ (Department of Water Affairs, 2013) which is decreased from natural conditions by water abstraction in the catchment, mainly through the Hazelmere dam located approximately 20 km from the estuary with a full storage capacity of 37.2 Mm^3^ (eThekwini Municipality, 2019). Outflows from wastewater treatment works provide some additional flow (ca 5 Mm^3^year^-1^, eThekwini Municipality) to the portion of river below the Hazelmere dam. The estuary itself is ca 1.5 km long (Begg, 1978). Depth can vary considerably between open and closed mouth conditions from ca 0.5 m to >2 m, respectively (this study).

**Figure 1:**
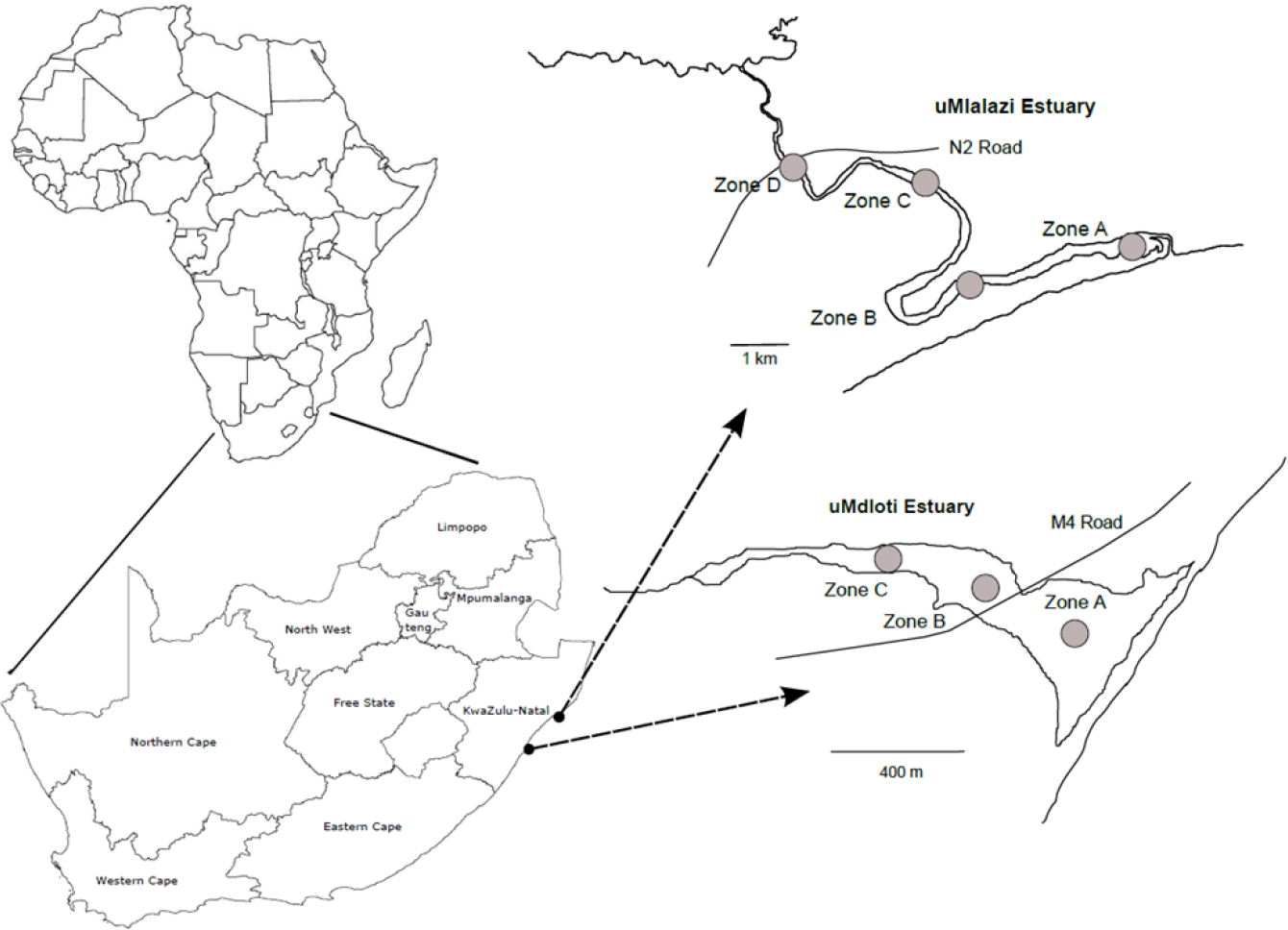
Maps of the uMlalazi and uMdloti estuaries, their estuarine zones and their location in South Africa. Grey dots denote sampling sites.

The predominantly open uMlalazi Estuary (31°49’22.8971”, 28°56’40.995”) on the northern part of the South African east coast is in a near natural condition. The flow is somewhat modified and the MAR is presently estimated at 124.6 Mm^3^ (van Niekerk et al. 2019). This estuary is of national biodiversity priority and an IBA (Important Bird and Biodiversity Area (BirdLife International; van Niekerk et al. 2019). It is about 11 km long (Begg, 1978), closes only under prolonged drought conditions, and has a maximum depth of 1 to 2.5 m along its length when open (DWS 2015).

Several environmental variables that are commonly used to investigate the distribution of biota in South African estuaries were measured during drought (2014-2016) and non-drought periods (2019- 2020). These include salinity, temperature (°C), dissolved oxygen concentration (mg/L) and pH measured with a YSI multiprobe (6600). Chlorophyll-a concentration was measured for total phytoplankton, as well as for three different size fractions of micro- (>20 µm), nano- (2-20 µm) and picophytoplankton. The picophytoplankton size fraction was filtered on 1.2 µm filters during drought and on 0.7 µm filters during non-drought conditions. Total phytoplankton filtered both on 1.2 and 0.7 µm filters showed a linear relationship (R = 0.95, p <0.001, n = 181) at a nearly perfect 1:1 line fit, which led us to assume that the density distributions of the picophytoplankton size fraction was not considerably influenced by the difference in filter size. Chl-a was extracted in 90% acetone at 4°C for 48 hours in the dark, and samples read on a Turner Designs 10-AU and Trilogy fluorometer. A significant linear relationship (R = 0.84, p < 0.001, n=176) was derived for readings of the same samples on both fluorometers which was used to derive chl-a values when one fluorometer was not available. All parameters were measured subsurface at ca 50 cm depth during daylight hours at neap tide, and the four physico-chemical parameters additionally near the bottom of the watercolumn.

Although nutrient concentrations are important in the context of primary producers, these were not included in the data analysis here due to a lack of data for the 2019/20 non-drought period. Sampling stations represent the lower, middle and upper estuarine reaches. Three stations were sampled in the comparatively shorter uMdloti Estuary (representing estuarine zone A, B and C; van Niekerk et al. 2019), and four in the longer uMlalazi Estuary (representing zones A, B, C and D; DWS 2015).

The entire dataset consists of bi-weekly measurements from June 2014 to May 2015 (drought period), and monthly measurements from June 2015 to June 2016 (drought period) as well as from June 2019 to March 2020 (non-drought period) (Fig 2). The 2019/2020 sampling series was cut short due to the COVID-19 pandemic. Density plots of the drought and non-drought period data were constructed and analysed to investigate the differences and similarities in their range and variability over time and space. To eliminate bias of the data representation in density plots and their statistical comparisons towards more, or less intensively measured time periods, the dataset was standardised to include the same number of time steps in the drought and non-drought periods. The common denominator for each drought and non-drought year was 10 time steps each. To consider responses in dry and wet seasons separately which are distinct in the climatic region in question (Dieppois et al., 2016; Roffe et al., 2021), the dataset is furthermore divided to represent dry (June to September) and wet (October to March) seasons. The spatial dimension is represented by gradients from the lower to the upper estuarine reaches. To compare the range and variability of the distributions, the analysis was conducted in two steps. First, the density distributions were statistically compared using a Kolmogorov-Smirnov test with a bootstrap hypothesis test of equality, through the R package ‘sm’ (Bowman and Azzalini, 1997). Here, the resampled mean differences are compared to the mean differences of our samples.

**Figure 2:**
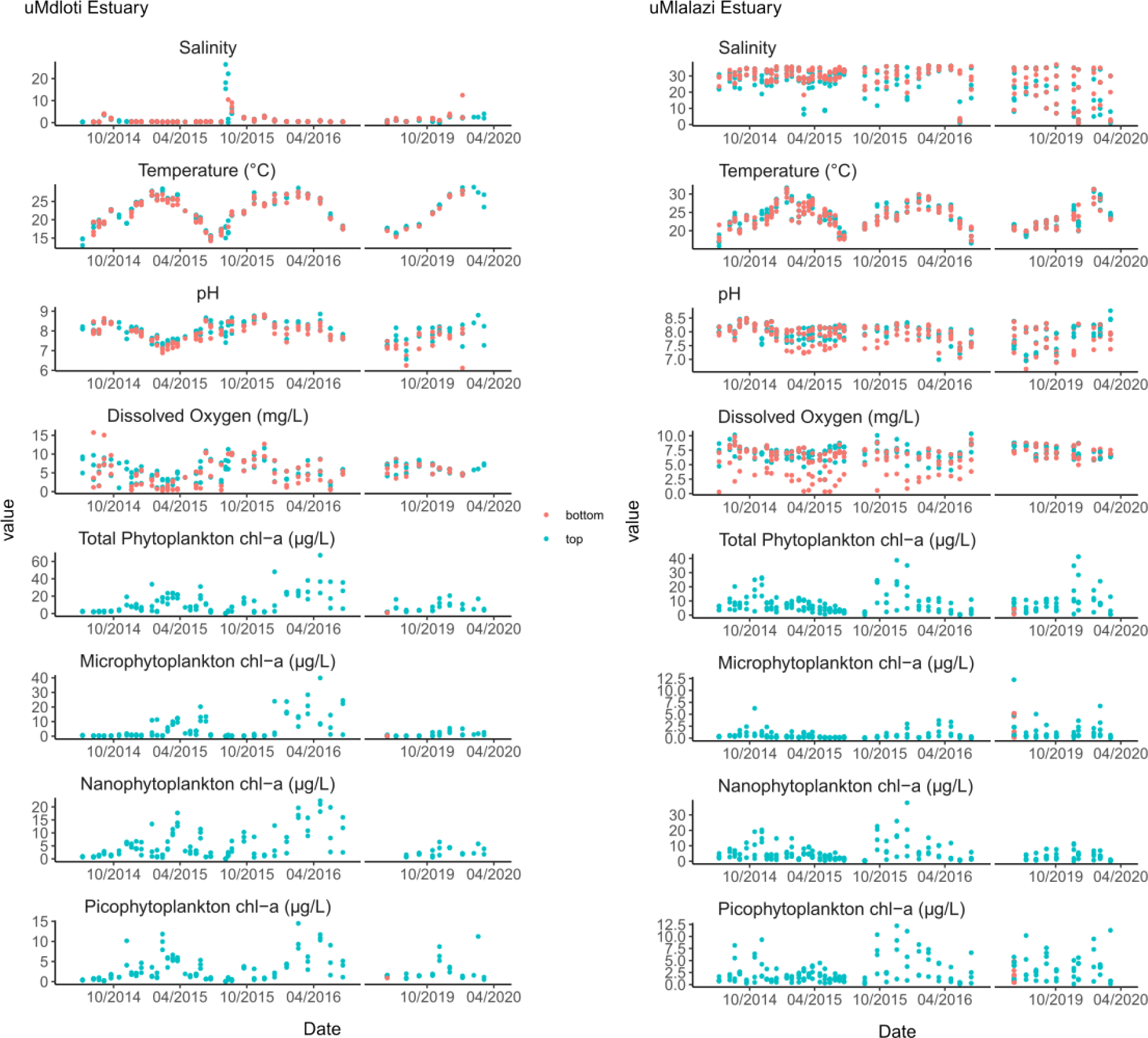
Physico-chemical parameters and phytoplankton biomass measured in drought (2014-2016) and non-drought (2019/20) periods in the uMdloti Estuary (left) and uMlalazi Estuary (right). Months on the x-axis show the start of the wet season (October) and start of the dry season (April).

Secondly, to evaluate both the extent of the divergence between two density distributions, and whether the divergence occurs in the lower, middle or upper value range, a Kullback-Leibler measure was calculated (Eq 1) using the function KLD in the R package ‘LaplacesDemon’ (Statisticat 2016):

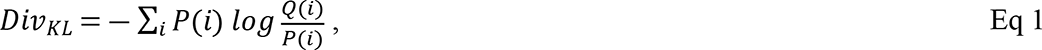

where *P* and *Q* refer to two different distributions that are compared, and *Q(i)* and *P(i)* are the density estimates at point *i* of the respective distributions. The results are presented as a sum of 512 point to point comparisons of the density distributions, and as plots of all 512 individual KL divergences to visualise whether the divergence occurs in the lower, middle or higher value range. This provides information on whether the drought and non-drought periods diverge more for their higher, or lower, for instance temperature range values.

The results for each estuary are presented separately, as the focus was on a drought/non-drought period comparison, rather than on comparing estuaries. Commonalities and distinct differences that arise from their different estuary type (temporarily closed, predominantly open), are however pointed out and discussed.

## Results

### Temporal trends

Several clear differences became apparent between drought and non-drought periods for the physico- chemical parameters and phytoplankton biomass. The trends are present in both wet and dry seasons (Fig 2, 3), in vertical stratification (Fig 2) and in spatial gradients between the lower and upper estuarine reaches (Fig 4, Table 1). Of the four physico-chemical parameters, the difference in salinity and dissolved oxygen (DO) concentration ranges and variability during drought and non-drought periods is the visually most striking for both estuaries (Fig 3). In LTCE uMdloti, salinities were consistently near zero during the drought years (2014-2016) except for one breach in 2015 that caused tidal intrusion and thus higher salinities for a very limited amount of time (Fig 2). The salinity during the non-drought period was also low, but with fewer measurements near zero especially during the wet season (Fig 2, 3). Its environment in terms of DO concentrations was highly variable during drought years, ranging from 0.24 – 12.7 mg/L, with a greater prevalence of low and hypoxic conditions throughout the estuary during the wet season, particularly in the first drought year (Fig. 2, 3). In contrast, DO was more stable, and more favourable during non-drought conditions when both very low (< 3mg/L) and very high (> 10 mg/L) concentrations were virtually absent and a narrower density peak settles near 6-7 mg/L in both the wet and dry season (Fig 3). A similar improvement in the stability of the DO environment was evident in the PrOE uMlalazi where density peaks are apparent between ca 7 and 8 mg/L during non-drought conditions (Fig 3). Here, low concentrations are absent especially from the bottom environment (Fig 2), important for benthic biota. Salinity on the other hand shows an opposite pattern to that in the uMdloti Estuary as in a predominantly open estuary increased riverine inflow lowers salinities in the upper reaches while tidal intrusion maintains high salinities in the lower reaches (Fig. 2, 3). Thus, regardless of season, higher freshwater flows in the non-drought year maintained typical spatial salinity gradients across the estuary, whereas low flow conditions during drought resulted in reduced salinity variation with poly- to euhaline salinities throughout. The estuary was overall cooler during the non-drought period (Fig 3). In both estuaries, the temperature environment was less variable during the non-drought period’s dry season and the high temperatures measured during the drought period were absent (Fig 3). pH was predominantly above 7 during the drought period, and featured lower values during the non-drought’s dry season resulting in a more variable pH environment especially during the dry season (Fig 2, 3).

**Figure 3:**
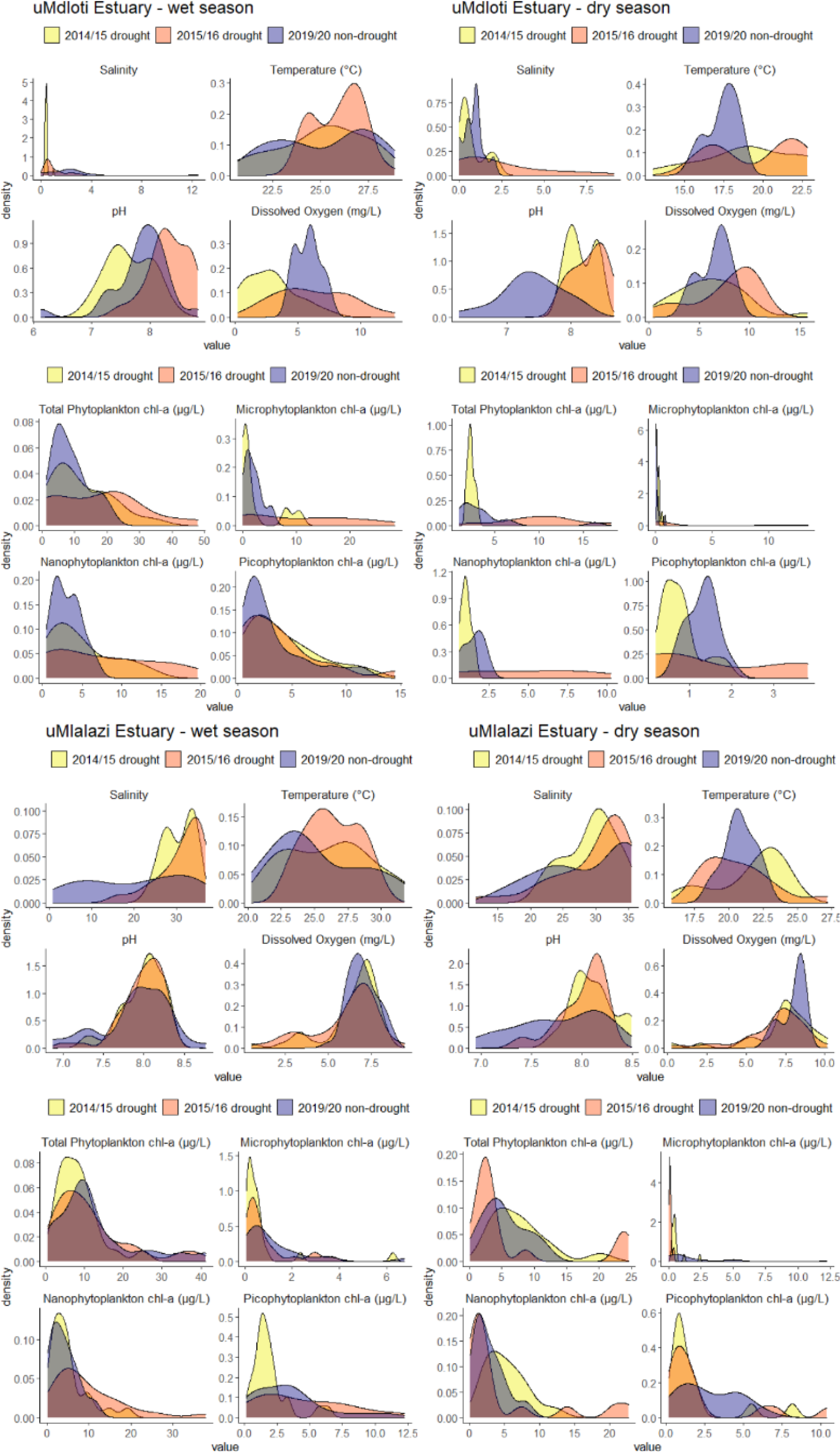
Density (y-axes) distributions of salinity, temperature, pH, dissolved oxygen concentration and phytoplankton chlorophyll-a concentrations (x-axes) during wet and dry seasons in the drought and non-drought years in the uMdloti and uMlalazi estuaries.

**Figure 4:**
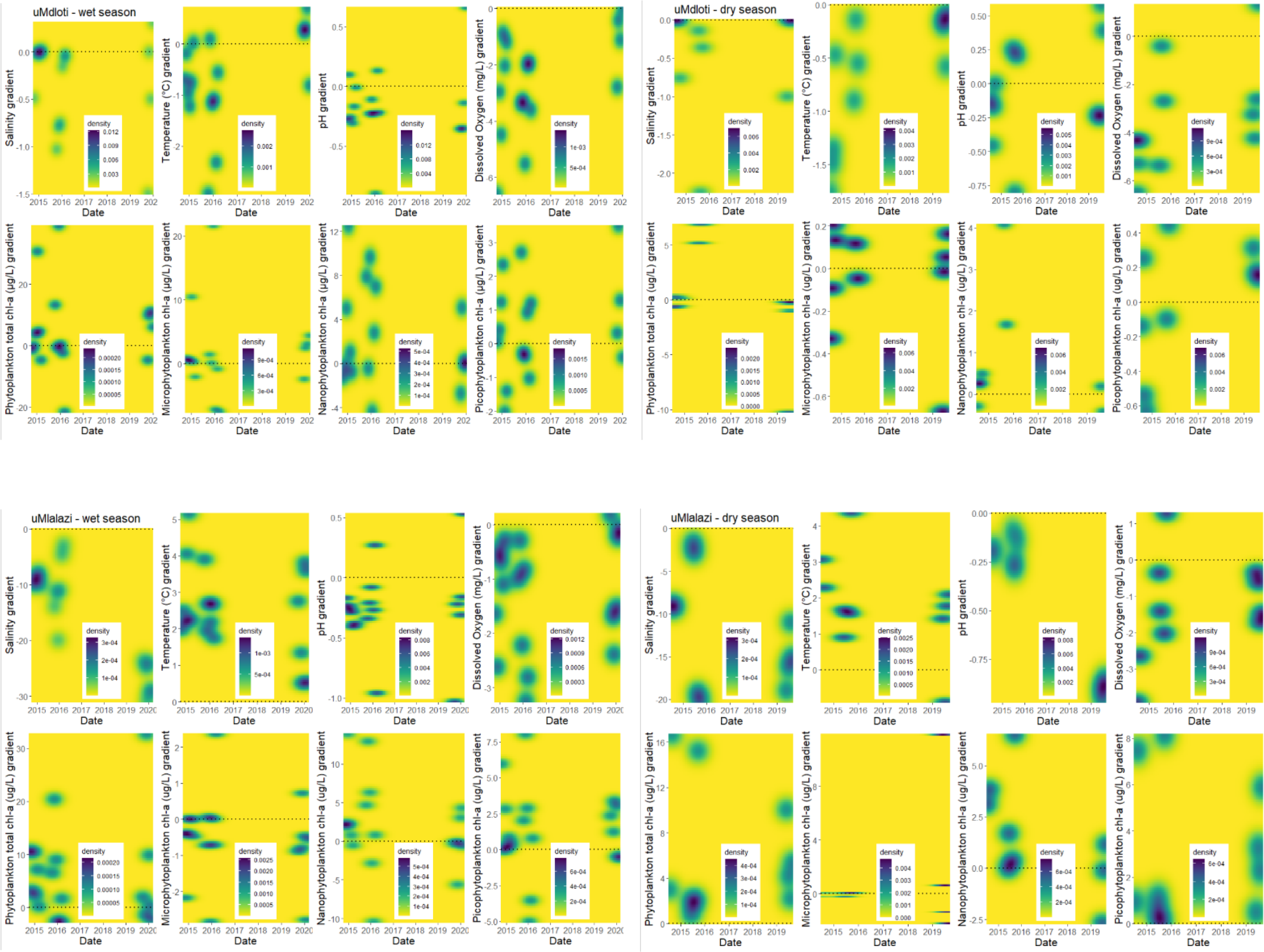
Density distributions of the spatial gradient between the upper and lower estuarine reaches (upper minus lower reach value) for salinity, temperature, pH, dissolved oxygen concentration and phytoplankton biomass during the wet and dry seasons of the drought (2014-2016) and non-drought (2019/20) years in the uMdloti and uMlalazi estuaries. The horizontal line denotes the position of no spatial gradient, points above the line (positive values) indicate a positive gradient from the lower to the upper reaches, and points below the line (negative values) a negative gradient from the lower to the upper reaches.

**Table 1:**
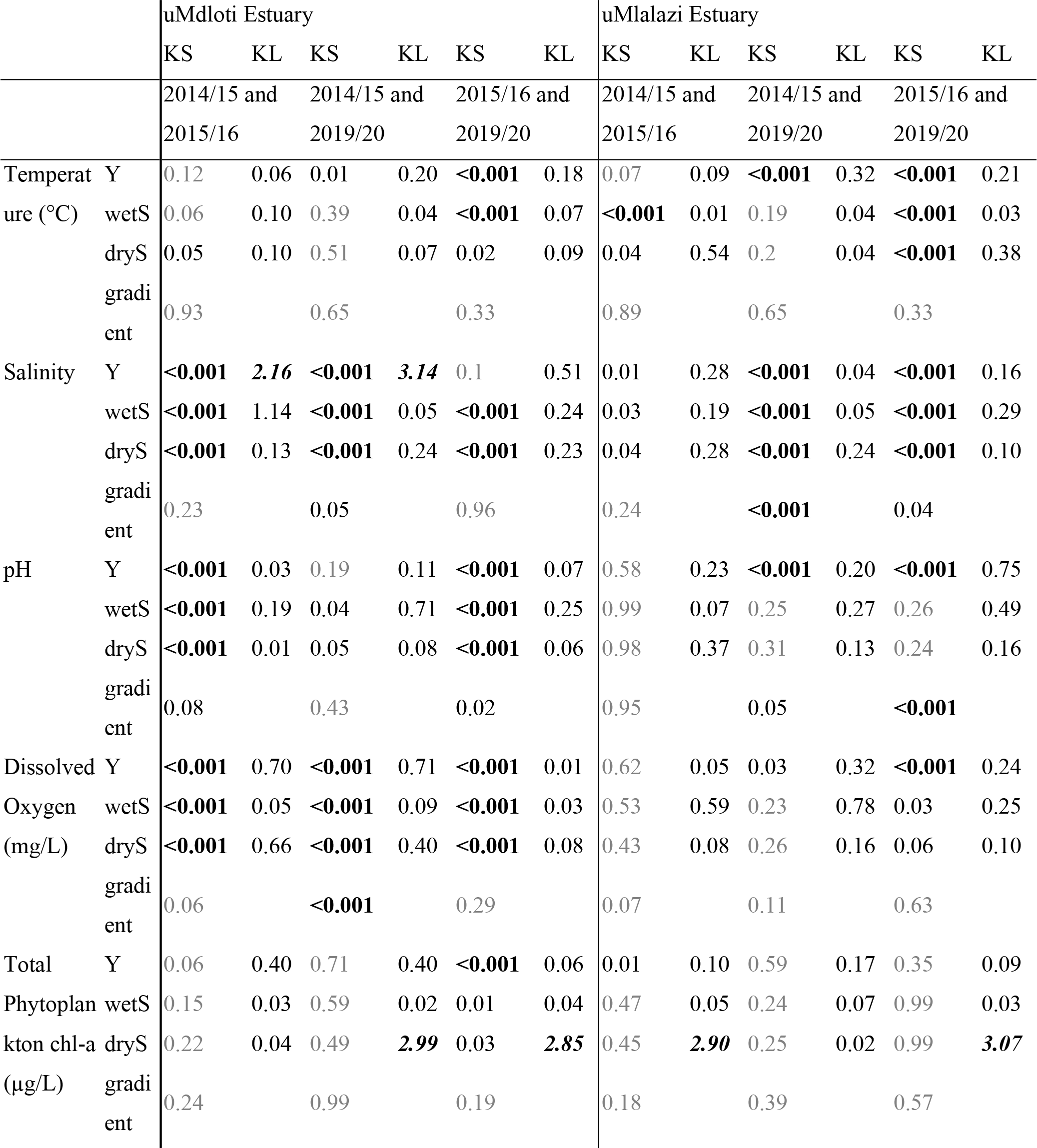
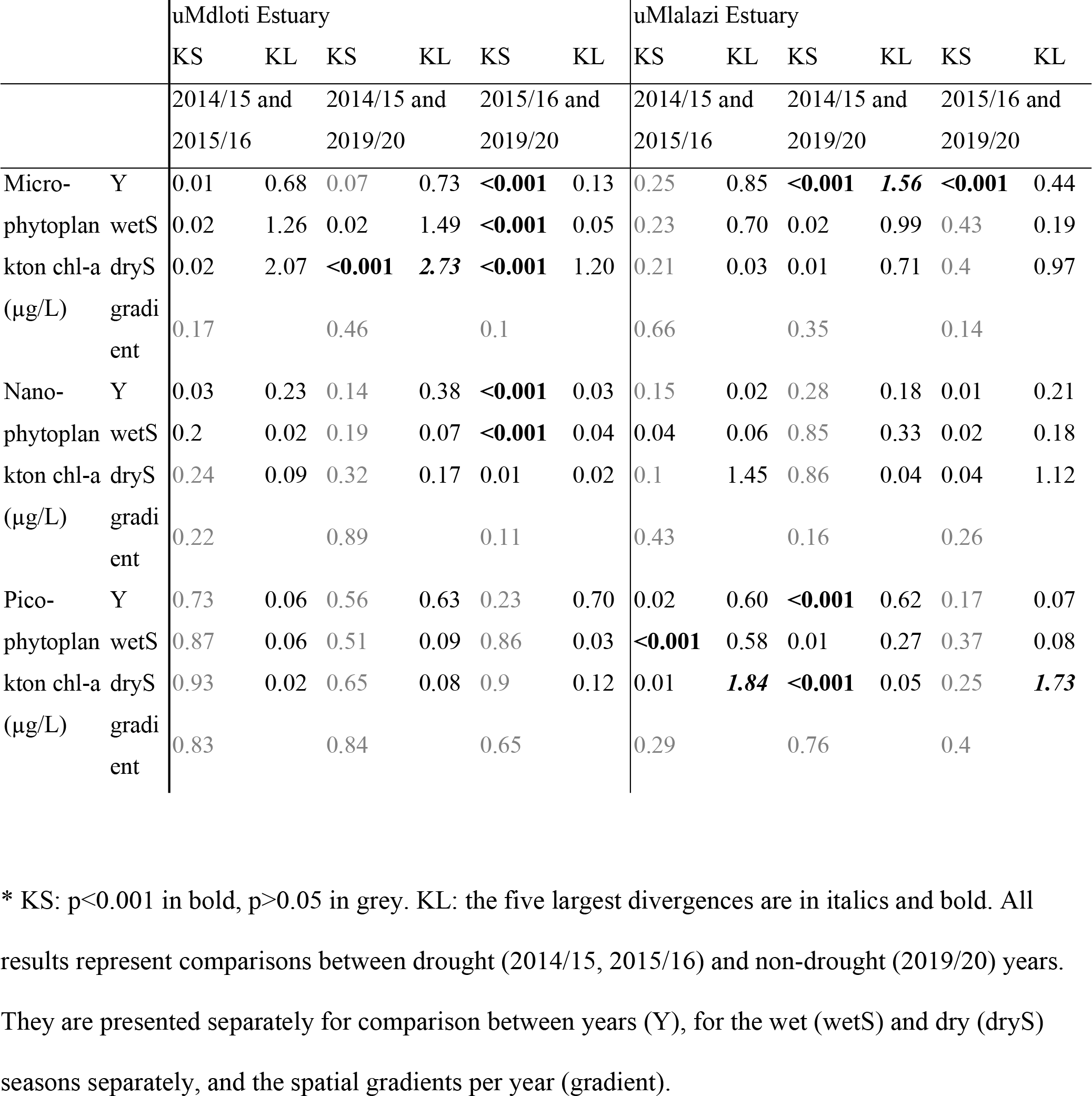
KS: Results (p-values) of the Kolmogorov-Smirnov permutation test of density distributions. KL: Kullback-Leibler divergence of density distributions. *

The variation in environmental parameters between drought and non-drought periods is partially reflected in the phytoplankton chlorophyll-a concentrations. These were generally lower during the non-drought period in the LTCE uMdloti (Fig 2). Peak phytoplankton chl-a biomass during drought ranged from 20-71 μg/L with higher values in the second year (2015/2016), whereas non-drought values did not exceed 20 μg/L. During spring and summer, chl-a concentrations in both estuaries increased, representing a phytoplankton growth period (Fig 2). Interestingly during the drought periods, chl-a for all phytoplankton size fractions in uMdloti peaked in a later season in summer/autumn (March, April) than during the non-drought period when chl-a concentrations peaked earlier in spring/summer (November, December). In the PrOE uMlalazi on the other hand, chl-a peaks in late spring were the norm during drought. During non-drought conditions, the clearest pattern comes from the total phytoplankton rather than from its size fractions, and showed a chl-a peak in summer (Fig 2). The combination of all three phytoplankton size fractions caused the pattern of total phytoplankton chl-a in both estuaries, but was driven predominantly by nano- and microphytoplankton in the LTCE uMdloti and by nano- and picophytoplankton in the PrOE uMalalzi (Fig 2). The total phytoplankton chl-a concentration may have influenced DO concentrations and pH, appearing as a contrary pattern over time that was more prominent in the uMdloti Estuary (Fig 2). Pearson’s correlations between total phytoplankton chl-a and DO concentration in the uMdloti were significant (R=-0.37, p< 0.001, n=190) only for the drought years, and between phytoplankton chl-a and pH in contrasting directions during drought (R=-0.42. p=<0.001, n=180) and non-drought (R=0.4, p=0.003, n=104) periods. In the PrOE uMalalzi on the other hand, only one fairly weak correlation was significant between phytoplankton chl-a and DO concentration (R=0.25, p=0.01, n=220).

Largest ranges of chl-a concentrations are evident during the drought period (Fig 3). In both drought and non-drought periods, the dry seasons showed smaller chl-a ranges and smaller variability (higher density peaks) in both estuaries. The response of phytoplankton chl-a to drought is thus different in terms of range of chl-a concentrations, and how often certain concentrations are reached (frequency). In addition, it is of importance whether measurements are taken during the dry or wet season when gauging the response. Statistically, in uMdloti the chl-a concentrations differed more often between the second drought year and the non-drought year, and most significant differences were caused by the microphytoplankton size fraction (Table 1). The picophytoplankton size fraction was the most different in uMlalazi and overall the first drought year was most often different from the non-drought year (Table 1).

### Divergence between drought and non-drought periods

To gauge in what way the dry and wet seasons during the drought and non-drought periods differ from one another, a Kullback-Leibler (KL) divergence measure was calculated in addition to statistically comparing the density curves. The statistical comparison renders a significant difference in cases where there is a consistent difference between density curves, or single large enough events during one period and not the other. The KL divergence measures a difference as the sum of divergences of all 512 point comparisons. The divergence therefore could yield a higher value due to single large rather than due to consistent small differences between the density curves. Therefore, highly statistically different comparisons do not necessarily conform with large KL values in Table 1. Still, both the statistical KS test result and the KL divergence are a summary measure of differences between two curves. For estuarine biota however it is important whether the lower, middle or higher value ranges differ. For instance, if the divergence for DO concentrations is apparent from the lower value range, this means that in one time period the habitat is more suitable than in the other, independent of a summary statistic concerning the entire value range.

Lower and middle range salinities diverged in the uMdloti more than the high salinity range values, which were virtually absent during both periods. The divergences were either due to the single breach in the 2015 dry season, or due to drought in the wet season particularly in 2014/2015 when freshwater conditions prevailed (< 0.5) (Fig 5). A shift towards lower salinities in the PrOE uMlalazi, especially during the wet season, is reflected in the divergences of the salinity density distributions in the higher and lower value ranges. The divergences follow from fewer higher and lower salinity measurements, whereas the middle range values were commonly encountered during both periods (Fig 5). DO concentrations diverged mostly in the middle and higher range values between drought and non-drought periods in the uMdloti Estuary (Fig 5). Even that both higher and lower range values are absent during the non-drought period, the especially high values during the first drought year’s dry season causes the difference in the higher range values. The lower range DO concentration measurements in the uMlalazi Estuary especially during the drought period’s dry season is apparent in the divergences. Generally the divergence peaks were larger for chl-a concentrations than for the physico-chemical parameters. The patterns are variable with respect to chl-a concentration divergences between drought and non-drought periods, and between wet and dry seasons, and reflect the different timing of the chl-a peaks, which in themselves show high variability in chl-a concentrations. The divergences were most prominent for the lower and middle range values rather than higher range values for both estuaries, and larger during the dry season (Fig 5). Of the phytoplankton size fractions, the microphytoplankton showed the largest divergence peaks in both estuaries. The information on divergences illustrated that distinct value ranges are affected differently rather than the entire value range uniformly.

**Figure 5:**
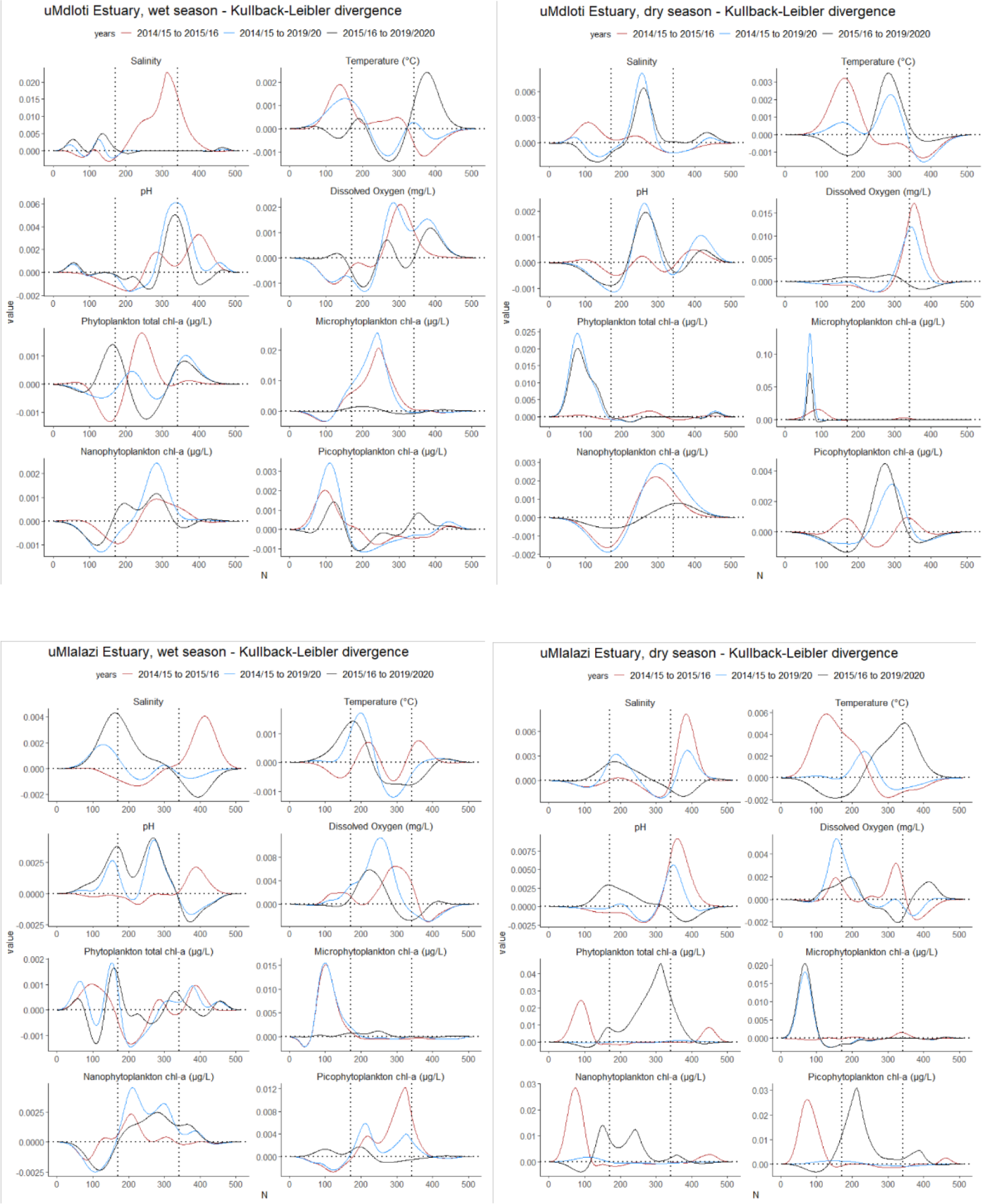
KL divergence (y-axis) of all variables for uMdloti and uMlalazi Estuary during wet and dry seasons for the drought and non-drought years. Compared are ca 500 points (N, x-axis) of the density distributions in Figure 3. The sum of all 500 points are represented as KL values in Table 1.

### Differences in spatial gradients

Lastly, we wanted to know whether there were also differences in estuarine axial gradients, i.e. whether the habitat niches expand or contract from the upper to the lower reaches, or whether the gradients remain the same? In the LTCE uMdloti, salinity gradients were the only ones to expand slightly from the drought to the non-drought period, whereas temperature, pH and DO concentration gradients contracted, during both wet and dry seasons (Fig 5, Table 1). In uMlalazi Estuary, salinity and pH gradients expanded, and temperature and DO concentration gradients contracted from the drought to the non-drought period (Fig 5, Table 1). When present, salinity, pH and DO concentration gradients were negative, indicating decreasing values from the lower to the upper reaches of the estuary. The temperature gradient during drought years in the uMdloti was negative, while a positive gradient was evident in the uMlalazi. The axial gradient of chl-a concentration mostly contracted during the non-drought period, but increased occasionally for total and for microphytoplankton chl-a in both estuaries (Fig 5). Thus, the environmental conditions in both estuaries not only showed fluctuations at individual estuarine reaches, but during non-drought conditions also provided (1) a larger range of habitat niches in terms of salinity gradients from the upper to the lower reaches, and (2) at the same time less variability and thus a more stable environment for most other parameters investigated.

The horizontal dotted line denotes zero divergence. The two vertical dotted lines separate the lower from the middle, and the middle from the higher range values.

## Discussion

A projected increase in ecological and agricultural droughts for South Africa (IPCC 2021) may lead to more frequent occurrences of environmental conditions such as those described in this study for the drought period. Together with anthropogenic activities that include abstracting water from estuarine catchments for agricultural, industrial and urban use, it is likely that future estuaries function differently compared to the present. Indeed, during the past decades, all South African estuaries have become, at least to some extent, threatened by flow modification (dams, water abstractions), with 20% severely threatened. This is a result of the abstraction of more than a third of the freshwater that used to reach the coast (van Niekerk et al. 2019). The combined pressures of anthropogenic flow modification and a change in climate may exacerbate the occurrence and duration of drought and its related effects on estuarine environments.

This study provides knowledge specifically on estuarine response to an El Niño induced drought. We discovered that environmental conditions had changed significantly between the drought and non-drought periods investigated. Drought conditions restricted the available salinity niches that are typical of estuaries, evident in the absence of higher salinity regions in the LTCE (uMdloti) and lower salinity regions in the PrOE (uMlalazi). When the mouth is closed, the uMdloti Estuary slowly turns fresh due to its perched nature and low but present freshwater input that partially arises from wastewater treatment work outflow (Brooker and Scharler, 2020). These shifts were not only apparent in the case of salinity, which is perhaps the most obvious parameter to change. A wider range of DO concentrations (including hypoxic values) and phytoplankton chl-a concentrations were measured during drought conditions. Phytoplankton chl-a concentrations are most probably not the only biota influencing DO concentrations in the uMdloti, as Pearson’s correlations between chl-a and DO concentrations for the dataset of the entire study period (R=-0.29, p<0.001, n=284) were statistically significant but not very strong. Benthic microalgae can outperform phytoplankton in terms of biomass, but have a record of a much lower production in uMdloti Estuary compared to that of phytoplankton (Anandraj et al. 2007). During our study at least in the first year, the mean estuarine microphytobenthos chl-a concentrations showed an opposite pattern to mean estuarine phytoplankton chl-a concentration and the DO concentrations are somewhat higher when microphytobenthos biomass is high (Supplementary material, Fig A). The relationship is however not statistically significant. DO concentrations could therefore be a reflection of both pelagic and benthic microalgae activity, that of filamentous macroalgae which bloom on occasion (this study, Omarjee et al., 2020), and in addition be influenced by the organic and bacteria load from the wastewater treatment works into the system.

Occasional blooms of the water hyacinth, *Eichhornia crassipes*, observed during the drought period further compete for nutrients and block out light. Therefore, although previously both riverflow gauged below the dam and the wastewater treatment work flows were shown to significantly influence phytoplankton chl-a concentrations (Brooker and Scharler 2020), the exact contribution and diel cycles of production and respiration of various primary producers and concomitant changes in DO concentration and pH in the estuary remain to be investigated. What can be concluded, however, is that both phytoplankton chl-a concentrations and the oxygen concentration environment are more variable with higher frequencies at extreme conditions during prolonged closed mouth periods during drought. A non-drought condition with more frequent mouth openings not only prevents hypoxia, but also facilitates a move towards a more stable and a more amenable oxygen environment. The mechanisms are tidal flushing, shorter residence time of water in the estuary, prevention of build-up of primary producer blooms (Taljaard et al. 2009), dilution of bacteria concentrations and more frequent replacement of deoxygenated water. Salinity and DO concentration measurements of the past confirm the special conditions during the drought period in the uMdloti Estuary of this study. During 1980/81, a non-drought period featured several mouth breaches with high DO saturation (> 75% at surface) and variable salinities (0-30) throughout (Blaber et al., 1984). About 20 years later in 2002/2003, over one year the salinity range was measured at 0-24 (Thomas et al., 2005), and DO concentrations at 0.8-11.9 mg/L (Kibirige et al., 2006). During these past studies, the estuarine inlet opened more frequently compared to the drought period of our study. The range of total phytoplankton chl-a concentrations during our study was overall smaller compared to 20 years ago (2002/2003) which is reported at 0.8 to 96 (Thomas et al., 2005) and 0.8 to 111 (Kibirige et al. 2007) mg chl-a/m^3^, even that input from wastewater treatment works did not decrease since then, but the frequency of inlet closure did, a period where higher chl-a biomass is often observed. During our study, the drought period was an extreme for salinity, DO and phytoplankton chl-a concentrations compared to the non-drought period.

In the predominantly open uMlalazi estuary on the other hand, the drought condition restricted the availability of the lower salinity niches through lower riverine discharge and unhindered marine intrusion that resulted in poly- and euhaline salinities throughout the estuary. Thus, the physical extent of habitat increased in the estuary during drought conditions for marine species. Estuaries with unhindered marine intrusions during drought periods elsewhere show an increase in marine species (Marques et al. 2014; Hallett et al. 2017). In the uMalalzi, the lower dissolved oxygen concentrations disappeared during the non-drought year – a distinct advantage for heterotrophic biota inhabiting the watercolumn and sediment. During the non-drought period, an additional station was sampled at the head of the estuary, which was inaccessible by boat during the drought period because of low water levels, due to build-up of sediment. At this station, salinity ranged between 5 and 20, and DO concentrations between 5.7 and 7.4 mg/L, which both lie within the range of the other estuarine stations. One and two decades earlier from our non-drought period, even wetter years produced salinity gradients of 0-35 (1999/2000: Mabaso, 2002; 2010/11: Ortega-Cisneros et al., 2014, Scharler unpub. data) and DO concentrations of 2.5 and 11 mg/L (Mabaso, 2002) and 0.7-10.3 mg/L, the latter with a mean of 6.7 and a median of 8.3 mg/L (Ortega-Cisneros et al., 2014; Scharler, unpub data). Evidently, the frequency of very low DO concentrations and high salinities was overall more pronounced during the drought years of the present study.

Are drought and non-drought periods simply extensions of naturally occurring dry and wet seasons in terms of estuarine environmental conditions? During different seasons, the environment changes for both temporarily closed and predominantly open estuaries, and their variability with respect to catchment land-use, freshwater availability and climate change responses provide different challenges to resident estuarine species and those using estuaries during part of their life cycle. Seasonal variability driven by temperature and light is compounded by freshwater inflow and mouth closures to which biota are adjusted by releasing larvae or recruit in and out of the estuary only during distinct times of the year that coincide with open inlets under natural conditions (Emmerson, 1994; Bell et al., 2001). Environmental parameters previously identified to influence the distribution of certain species in certain estuaries may react to different extents to the various global change drivers. The results vary whether the changes are short or long term and thus another hurdle to upscaling effects from dry seasons to droughts is the duration of each. A drought usually lasts more than one season, and effects of droughts may have longer-term impacts on estuarine populations beyond the end of a drought (Wetz et al., 2011). Patterns apparent from the wet and dry season therefore are not directly scalable to drought and non-drought periods. Drought can even have a larger impact on estuarine water quality compared to different land-use types (Elsdon et al., 2009), and are more pronounced in estuaries that are subjected to artificial freshwater abstraction from their catchments. For instance, predominantly open estuaries that are already typically euhaline throughout become hypersaline in the upper reaches during low flow periods, such as the temperate Kromme Estuary in South Africa (Scharler and Baird 2000). Similar to the uMlalazi, the effects of a prolonged drought on the Gamtoos Estuary resulted in the short-term closure of the mouth, with further effects of homogeneous polyhaline salinities, a shift towards eutrophic conditions, hypoxia in bottom waters, but restoration of typical estuarine gradients when the mouth reopened (Lemley and Adams, 2020). Since estuaries are largely individual in their responses due to differences in land-use, morphology, freshwater availability, and others, disentangling these effects may yield different results for different estuaries (Chuwen et al. 2009; Elsdon et al. 2009).

Arid regions and those with seasonal or periodic freshwater scarcity occur across the globe. Hence, impacts on estuarine biota of reduced riverine flow during drought have been documented for a wide range of biota, but mostly concentrated on fish. Still, all components of the food web are affected when community changes such as species diversity and biomass are induced by drought. Depending on the estuary, phytoplankton biomass may be higher (Brazil: Barroso et al., 2018; USA: Cira et al., 2021) or lower (Brazil: Pereira et al., 2017; USA: Wetz et al., 2011) during drought. Higher biomass can be concomitant with lower biodiversity (Cira et al., 2021). Phytoplankton in estuarine systems have in the past more generally been pinpointed as variable and depending on the specific circumstance of annual climatology, and specific disturbances affecting the system (Cloern and Jassby 2010; Schallenberg et al. 2010). Our data are relatively short term, but we still detected a difference between disturbance (drought), between seasons (shift in phytoplankton chl-a peak) and in addition quantified the differential impact during wet and dry seasons, and which value ranges were more, or less impacted.

The impact received by a semi-urban estuary (LTCE uMdloti) is lacking in the PrOE uMlalazi, but also here we detected high variability. For both estuaries it remains to be seen from future studies whether the divergencies in chl-a always occur in the lower to middle value ranges, or whether always the same size fractions imprint their variability on that of total phytoplankton chl-a.

The variabilities of the changing food supply from trophic level 1 (e.g. phytoplankton) may affect the entire food web beyond physiological preferences of consumers. Invertebrates have been documented to be severely affected by drought, through changes in the estuarine environment. For instance, the hydroclimate has a profound influence on estuarine zooplankton in Portugal (Marques et al., 2014), and both temperature and dissolved oxygen concentrations affect invertebrates in the Thames estuary (Attrill and Power, 2000). In both the southern USA and southern Australia, species specific effects of drought on fish impact abundances either positively or negatively (Dolbeth et al., 2010; Wedderburn et al., 2012; Mickle et al., 2018). When drought induced closure of the estuarine inlet occurs, not only are the extent and variability of the physico-chemical environment, and community composition affected. Recruitment migrations into the estuary and back out to sea of larvae, juveniles and adults are restricted, which may even lead to temporal species extinctions (Carrasco et al. 2013; Whitfield 2021). Fishery declines have been linked to drought in eastern Australia (Gillson et al., 2009) and specifically to a drought induced lack of prawn recruitment in South Africa (Ayers et al., 2013).

## Conclusion

Although changes induced by drought can have such variable effects on estuaries and their biota, we can draw several conclusions from our study. Under more frequent or prolonged droughts, species inhabiting a typically perched LTC subtropical South African Estuary (uMdloti) may expect extremely low salinities, a higher frequency of higher temperatures, and more variable DO concentrations including hypoxic conditions. The phytoplankton component during drought years is more variable for all size fractions. The expected changes for biota during drought are a shift in biodiversity towards higher occurrence of freshwater species, and a loss of marine and estuarine species (Scharler et al. 2020), as well as limited recruitment due to a closed mouth. In the predominantly open uMlalazi Estuary, we expect salinities and temperatures restricted to higher value ranges, less variable pH and more variable DO concentrations during drought conditions. Phytoplankton chl-a concentrations will be higher and more variable. For other biota we can expect more marine and estuarine, and fewer freshwater species, and biotic exchange with the sea is most likely hindered only occasionally when the mouth closes. The marinisation of the predominantly open estuary conforms with predictions for south west Australian permanently open estuaries (Hallett et al., 2018). Temporarily closed estuaries to the contrary can have different trajectories from complete freshening to hypersalinity, depending on whether river flow ceases completely or not, their morphology, whether it is perched, whether the mouth is open or closed, and to different degrees on land-use of adjacent ecosystems.

## Acknowledgements

We would like to thank the following people for assistance in the field: Fru Azinwi Nche-Fambo, Thembeka Radebe, Dane Garvie, Kajal Lechman, Andile Nkosi, Dikarabo Rafedile, Gemma Gerber, Ben Brooker and Siyabonga Biyase. GG and DK also assisted in the data cleaning of the 2019/20 dataset. This work was financed by National Research Foundation Grants (Grant No: 93554 and 118608).

## Supplementary Material

Microphytobenthos samples for chlorophyll-a measurements were taken at each site in each estuary, concurrently with other samples and measurements as described in the methods section of the main text. Three replicates were sampled at each site using a corer with a diameter of 2 cm. The top 1 cm of the sediment core was added to 30 ml of 90% acetone, and extracted and analysed in the same way as the chl-a of phytoplankton described in the main text.

**Figure A:**
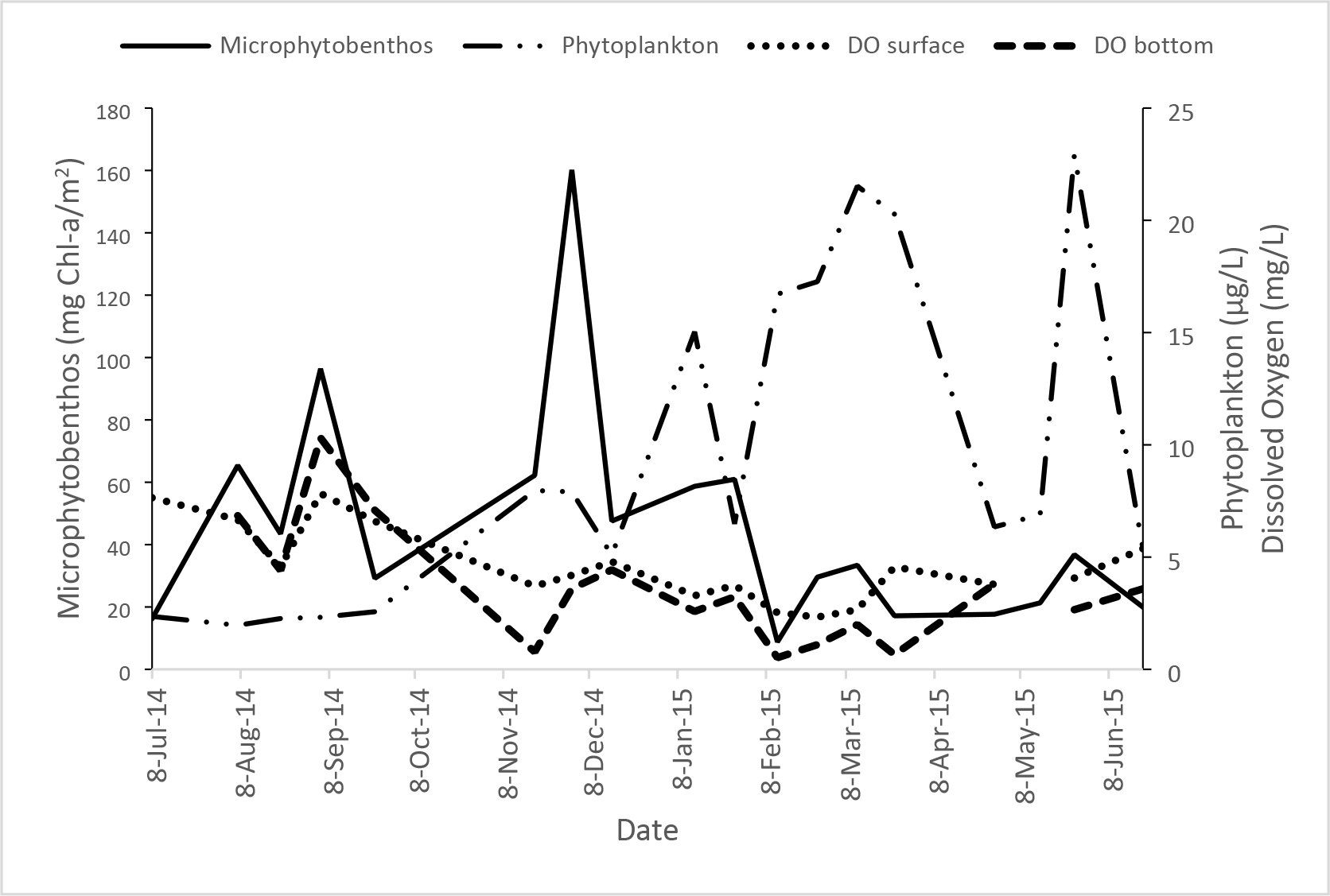
Mean estuarine phytoplankton and microphytobenthos chl-a concentrations, and dissolved oxygen concentrations (surface and bottom of watercolumn) over one year illustrate the potential contribution of benthic primary producers to watercolumn DO concentration in the uMdloti Estuary. Shown is the time period with the most complete dataset for both phytoplankton and microphytobenthos data over the study period (this study).

## Notes

### Competing Interest Statement

The authors have declared no competing interest.

